# Synthetic logic circuits using RNA aptamer against T7 RNA polymerase

**DOI:** 10.1101/008771

**Authors:** Jongmin Kim, Juan F. Quijano, Enoch Yeung, Richard M. Murray

## Abstract

Recent advances in nucleic acids engineering introduced several RNA-based regulatory components for synthetic gene circuits, expanding the toolsets to engineer organisms. In this work, we designed genetic circuits implementing an RNA aptamer previously described to have the capability of binding to the T7 RNA polymerase and inhibiting its activity *in vitro*. Using *in vitro* transcription assays, we first demonstrated the utility of the RNA aptamer in combination with programmable synthetic transcription networks. As a step to quickly assess the feasibility of aptamer functions *in vivo*, a cell-free expression system was used as a breadboard to emulate the *in vivo* conditions of *E. coli*. We tested the aptamer and its three sequence variants in the cell-free expression system, verifying the aptamer functionality in the cell-free testbed. *In vivo* expression of aptamer and its variants demonstrated control over GFP expression driven by T7 RNA polymerase with different response curves, indicating its ability to serve as building blocks for both logic circuits and transcriptional cascades. This work elucidates the potential of RNA-based regulators for cell programming with improved controllability leveraging the fast production and degradation time scales of RNA molecules.

## Introduction

Synthetic biology provides an engineering approach to the field of biology to (re)design tools and biological systems that can open up new possibilities in biotechnology and medicine [1, 2, 3, 4, 5]. To achieve this goal, such *de novo* biological systems need to be easily engineerable, ideally through utilizing molecular mechanisms that work much as the programming codes. The first synthetic gene networks, the toggle switch [6] and the repressilator [7], demonstrated the feasibility of programming computational tasks in living cells. Recent advances in synthetic biology further expanded the set of programs in biological organisms with novel functions including logic gates [8, 9, 10, 11, 12], cell classifiers [13], and analog signal processors [14]. Most synthetic gene circuits constructed to date relied on well-characterized protein components, which results in a challenge for scaling up synthetic circuitry with predictable dynamical behaviors due to the idiosyncratic nature of each protein component involved. Recent advances aim to address such limitations through increasing the number of orthogonal protein regulators [15] as well as protein regulators that rely on nucleic acids as guiding molecules [16, 17].

To address the component bottleneck in synthetic biology, RNA structures have also been utilized as important tools for programming gene expression inspired by the naturally occuring nucleic-acid-based regulatory motifs [18, 19, 20]. By virtue of their predictive interactions with other nucleic acids, proteins, and small molecules, many RNA-based regulatory circuits have been investigated and synthesized [21, 22, 23, 24, 25, 26, 27, 28, 29, 30, 31, 32]. Importantly, RNA aptamers possess the molecular recognition properties with specificity and affinity rivaling protein antibodies for exclusive ligands, allowing these molecules to form a useful set of components for synthetic circuitry in living cells. RNA aptamers are typically generated and selected *in vitro* starting from large random sequence libraries, and are subsequently optimized for high affinity binding to given ligands by a process known as SELEX (Systematic Evolution of Ligands by EXponential enrichment) [33, 34, 35, 36].

In a recent work, Ohuchi and co-workers [37] reported the *in vitro* evolution of RNA aptamers against T7 RNA polymerase (T7 RNAP) through SELEX. Although the detailed mechanism is not yet fully understood, it was shown that the RNA aptamers inhibited the association of T7 RNAP with T7 promoter DNA, thereby inhibiting the activity of RNAP in *in vitro* transcription assays. Utilizing the property of this aptamer, we implemented synthetic logic circuits in an *in vitro* transcription setting. As a step to quickly assess the aptamer functionality *in vivo*, a cell-free transcription-translation (TX-TL) system was employed as a ‘breadboard’ to emulate the *in vivo* conditions of *E. coli* for circuitry testing [38, 39]. Experimental results demonstrated that the T7 RNAP aptamer showed up to 8-and 5-fold inhibition in TX-TL system and in *E. coli* MGZ1 cells, respectively. Due to the wide-spread usage of T7 RNAP for *in vitro* and *in vivo* applications [40, 41, 42, 43], the T7 RNAP aptamer can be a useful addition to the regulatory toolkit for *in vitro* and *in vivo* synthetic biology.

## Materials and Methods

### Circuit construction and use of strains

Plasmid pAR1219 was obtained from Sigma-Aldrich. Plasmid pTara was provided by Matthew Bennett’s group at Rice University. The rest of the circuits were constructed using Gibson assembly [44]. Schematics for plasmids are shown in the main text and the full sequences and annotations are described in Supplementary Figure S1. Plasmids were transformed into strains via chemical transformation. *E. coli* MGZ1 strain (MG1655 containing the LacI-and TetR-overexpressing Z1 cassette) was used for inducible expression from the TetR-regulated promoter. Plasmids pdeGFP-T7apt-4A, pdeGFP-T7apt-4G, and pdeGFP-T7apt-4Δ were derived from the plasmid pdeGFP-T7apt via round-the-horn site-directed mutagenesis using 5′-phosphorylated primers for subsequent ligation. Plasmids containing the T7 RNAP aptamer and variants have an additional XhoI site between the aptamer and the terminator sequences (the functionality of the T7 RNAP aptamer was not affected by this feature). All of the plasmid sequences were confirmed by DNA sequencing.

### *In vitro* transcription reaction

The sequence of all DNA molecules and expected RNA transcript sequences were chosen to minimize the occurrence of alternative secondary structures using the DNA and RNA folding program NUPACK [45]. The DNA and RNA sequences used in this study are listed in Supplementary material. All DNA oligonucleotides were purchased from Integrated DNA Technologies (USA). The T7 RNA polymerase (Cellscript, Madison, WI, USA; #C-AS2607), T3 RNA polymerase (Promega, Madison, WI, USA; #P4024), 10*×* transcription buffer and thermostable inorganic pyrophosphatase (New England Biolabs, Ipswich, MA, USA; #B9012S, #M0296S), NTP (Epicentre, Madison, WI, USA; #RN02825) were purchased. Malachite green dye was purchased from Sigma (#M9015). Since pyrophosphatase is involved in regulating the byproduct inorganic pyrophosphate for our transcriptional circuits and is not directly involved in the dynamics, we do not call it an “essential enzyme” for the circuit dynamics.

The fluorescence was recorded every minute using a Fluorolog-3 spectrofluorometer (Jobin Yvon, Edison, NJ, USA). The excitation and emission for malachite green fluorescence were at 630 nm and 655 nm. DNA templates were annealed with 10% (v/v) 10*×* transcription buffer from 90*^◦^*C to 20*^◦^*C over 1 hour at 5 *µ*M concentrations. Transcription reactions were prepared by combining the annealed templates, 7.5 mM of each NTP, 24 mM MgCl_2_ (to balance salt concentrations due to increased NTP concentrations), 10% (v/v) 10*×* transcription buffer, and 25 *µ*M malachite green dye in a test tube. Transcription reactions for spectrofluorometer experiments were prepared as a total volume of 60 *µ*L in quartz cuvettes and enzymes (RNAP and PPase) were added and mixed after the baseline fluorescence was recorded for 10 min. Sample temperature was maintained at 29*^◦^*C using a 4-sample changer with a temperature-controlled water bath.

### Media, chemicals and other reagents

Strains were grown in LB liquid medium or on agar plates (1.5% agar), supplemented with carbenicillin (100 *µ*g/ml), and chloramphenicol (34 *µ*g/ml) as needed. IPTG induction was at 10 *µ*M unless otherwise noted. Anhydrotetracycline (aTc) induction experiments used different concentrations of aTc to achieve different levels of RNA aptamer expression as described in the figure captions.

### Plate reader analysis

For cell-free TX-TL experiments, GFP fluorescence was measured using a filter set with excitation wavelength at 488 nm and emission wavelength at 507 nm. All data were obtained using Perkin Elmer Victor X3 plate reader. Preparation of the cell-free TX-TL expression system was done according to previously described protocols [38, 39]. A single batch of *E. coli* extract was used consistently throughout the experiments to prevent variation from batch to batch. Samples were prepared by combining extract and buffer tubes freshly thawed from *−*80*^◦^*C, and purified plasmid DNA. Concentrations of plasmids pAR1219 and pdeGFP-T7apt and its variants were 0.8 nM for Figure 2. The concentrations of plasmids pAR1219, pT7-deGFP, and pTet-T7apt were 0.6 nM for Supplementary Figure S3. The reaction volume was 10 *µ*L in a 384-well plate (Nunc), and the number of replicates was two for each reaction. Sample temperature was maintained at 29*^◦^*C.

**Figure 2:**
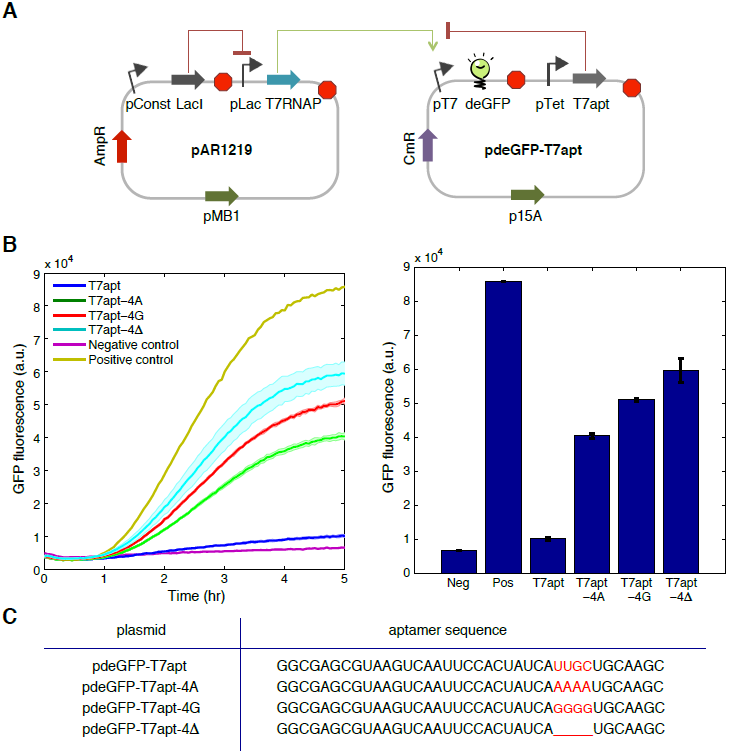
Characterization of T7 RNAP aptamer and its variants in the cell-free TX-TL system. (**A**) Two-plasmid system used for aptamer-mediated T7 RNAP inhibition test in the TX-TL cell-free extract. (**B**) The expression of T7 RNAP was induced by IPTG and the effect of expression of T7 RNAP aptamer and its variants was tested in TX-TL system. TX-TL results showed that deGFP expression was approximately 8-fold decreased in the presence of the correct aptamer; the fluorescence signals were much higher in the presence of modified aptamers indicating disrupted functionality. Negative control contained only pAR1219 plasmid and positive control contained pAR1219 and pT7-deGFP plasmids. The T7 RNAP aptamer production was not inhibited by TetR since the cell extract does not contain TetR protein. The fluorescence signal from deGFP was measured at 5 hrs for the bar plot. (**C**) The sequence of T7 RNAP aptamer and its variants. Red color highlights the modified consensus sequence domain.

For *in vivo* experiments, cells were grown in shaking liquid culture (LB media with carbenillin and chloramphenicol) at 37*^◦^*C overnight. Following this, the culture was diluted 1/200 fold for plate reader measurements. GFP fluorescence was measured using a monochromater with excitation wavelength at 485 nm and emission wavelength at 525 nm in Biotek Synergy H1 plate reader. Fluorescence data were normalized by OD600 values. The culture volume was 200 *µ*L in a 96-well plate (Nunc), and the number of replicates was three for each condition. Sample temperature was maintained at 37*^◦^*C.

### Flow cytometry analysis

Cells were grown in shaking liquid culture (LB media with carbenillin and chloramphenicol) at 37*^◦^*C overnight. Following this, the culture was diluted 1/100 fold and grown at 37*^◦^*C (LB media with antibiotics and 1% glucose and aTc) for two hours. Flow cytometry analysis used a Cell Lab Quanta SC MPL flow cytometer (Beckman Coulter). GFP was excited from a 488 nm laser and emission was measured from 517 nm longpass filter with a PMT setting of 5.0. For each sample, the GFP values of 50,000 cells were measured. Cells were then gated for GFP levels above background (2 a.u.). Data were analyzed using MATLAB and modal values were determined. The number of replicates was three for T7apt and the number of replicates were two for its variants T7apt-4A, T7apt-4G, and T7apt-4Δ.

## Results

### T7 RNAP aptamer for *in vitro* transcriptional circuits

First, we aim to demonstrate the utility of T7 RNAP aptamer for *in vitro* transcriptional circuits. Using the synthetic transcriptional switch as the regulatory motif and the aptamer for the chromophore malachite green (MG) as the output signal, we construct a logic circuit utilizing the property of T7 RNAP aptamer (Figure 1A). Among the several variants of T7 RNAP aptamers reported in Ohuchi et al. [37], our first *in vitro* assays utilized the aptamer T230-38. In this circuit, the two RNA polymerases, T3 RNAP and T7 RNAP, together with DNA activators serve as inputs and the fluorescence of MG aptamer serves as an output. Figure 1B illustrates the molecular reactions for this synthetic transcriptional circuit. An MG aptamer, MGapt, consists of a short RNA sequence whose central loop region serves as the binding pocket for MG. When MG is bound to the aptamer, it becomes highly fluorescent [46], thereby providing a convenient, real-time read-out for RNA expression. Previous *in vitro* synthetic transcriptional circuits utilized T7 RNAP [47, 48, 49, 50, 51]. However, since we aim to characterize an aptamer against T7 RNAP, a separate enzyme was required to drive the production of aptamer in order to avoid self-repression. Thus, we first verified the design principle for synthetic switches utilizing T3 RNAP using a template encoding MG aptamer with an incomplete promoter for T3 RNAP, pT3-MGapt; the production of MGapt was observed upon the addition of its cognate activator, input A2 (Supplementary Figure S2A). Utilizing the modular switch architecture where the output domain is physically separated from the input and promoter domains, the two templates were designed to transcribe different RNA outputs and to be recognized by two different RNA polymerases. (See Supplementary Method for sequences.) The two templates encoding T7apt and MGapt are hereby denoted as pT3-T7apt and pT7-MGapt. In this circuit, T3 RNAP recognizes pT3 promoter upon addition of the DNA activator A2, and induces transcription of T7apt; T7 RNAP recognizes pT7 promoter upon addition of the DNA activator A1, and induces transcription of MGapt.

**Figure 1:**
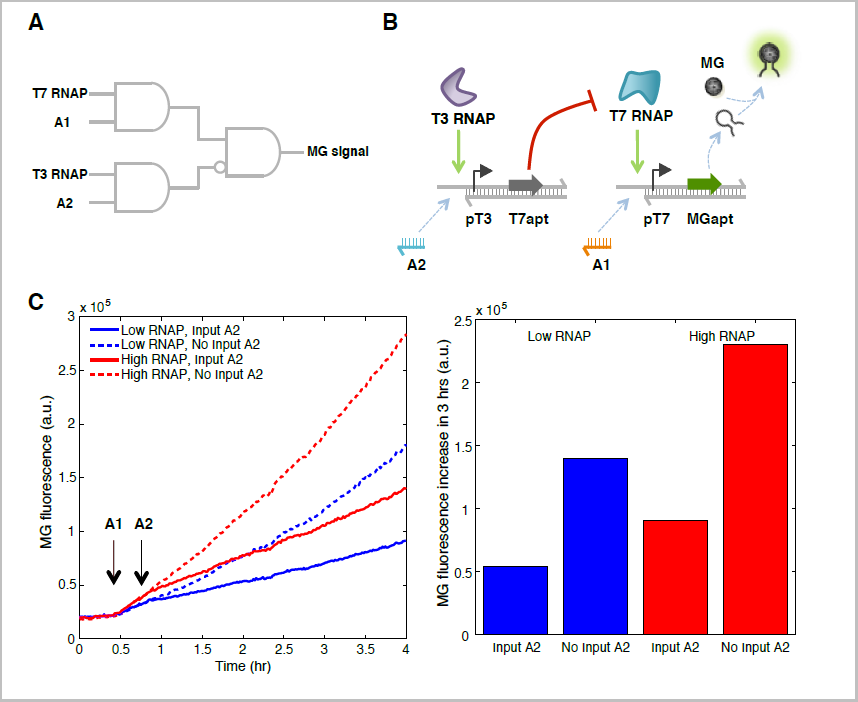
Characterization of T7 RNAP aptamer in *in vitro* transcription circuits. (**A**) Schematic representation of logic circuit. The two RNA polymerases and DNA activators serve as inputs and the fluorescence of malachite green aptamer serves as an ourput. (**B**) Detailed diagram of synthetic transcription circuit. Promoters pT3 and pT7 are to be recognized by T3 RNAP and T7 RNAP, respectively, upon addition of the corresponding DNA activator A2 and A1. The switch templates contain a promoter for RNA polymerase that lacks part of the double-stranded sequence; after addition of a single stranded DNA activator, the promoter is complete except for a nick, and thus, the transcription efficiency is high. When the MGapt is expressed, it can binds to the chemical compound malachite green, greatly increasing the fluorescence signal output. (**C**) Transcriptional assay of the circuit shown in (B). The two RNA polymerases, T3 RNAP and T7 RNAP, were added at 10 min, the input DNA activator A1 was introduced at 25 min and the input DNA activator A2 was added at 45 min as marked by black arrows. The T7apt was efficient in repressing the T7 RNAP activity; about 3-fold decrease in MGapt signal was observed over 3 hours in the presence of T7apt. ‘Low RNAP’ and ‘High RNAP’ refer to T7 RNAP concentrations of 20 and 40 nM, respectively.

Experimentally, we initially included both templates pT3-T7apt and pT7-MGapt, both T3 and T7 RNAP along with NTP fuel in the reaction mixture (Figure 1C). No MGapt expression was observed until the addition of DNA activator A1; upon the addition of input A1, the transcription of MGapt was immediately observed. Then, the DNA activator A2 was added for half of the samples, enabling transcription of T7apt in order to inhibit T7 RNAP-mediated MGapt expression (Figure 1C). The ‘input A2’ trajectories began to diverge from ‘no input A2’ trajectories about 10 min after the introduction of input A2 (Supplementary Figure S2B). The T7apt was efficient in repressing the T7 RNAP activity; about 3-fold decrease in MGapt signal was observed in the presence of T7apt. Further, we tested a modified T7apt termed T7apt-v2 where two bases comprising stem were switched and six additional nucleotides were attached at the 3′ end; T7apt-v2 was equally efficient in repressing the activity of T7 RNAP (Supplementary Figure S2C), indicating that the core structural motif of T7apt is sufficient for its activity.

### T7 RNAP aptamer functionality in cell-free TX-TL system

After observing the T7 RNAP aptamer functionality as a circuit element in the synthetic transcription system, we designed a new circuit for *in vivo* application (Figure 2A and Supplementary Figure S3A). Creating a functional *in vivo* biocircuit can be a lengthy process, involving multiple iterations of design, assembly, and test cycles. Similar challenges may arise for testing novel regulatory mechanisms such as the inhibition of T7 RNAP through aptamers. To streamline and expedite the design-build-test cycles, we chose to employ cell-free TX-TL system; TX-TL system is gaining interest as biomolecular breadboards that take advantage of shorter assembly and troubleshooting times by conducting reactions *in vitro* emulating *in vivo* conditions [38, 39]. Because requirements for propagating plasmids *in vivo* have been removed, fast iterations of the design-assembly-test cycle of a given circuit is possible using linear DNA constructs [52]. Further, RNA-based regulatory elements were recently shown to behave analogously in TX-TL system when compared to *in vivo*, increasing the utility of this platform [53].

In order to emulate the T7 RNAP aptamer functioning *in vivo*, we assayed the aptamer in the TX-TL system. Initially, a three plasmid system was tested (Supplementary Figure S3B): plasmid pAR1219 encoding T7 RNAP under pLac promoter along with LacI constitutively expressed, plasmid pT7-deGFP encoding deGFP (truncated eGFP to maximize fluorescence output [38]) under pT7 promoter, and plasmid pTet-T7apt encoding T7 RNAP aptamer under pTet promoter. For a negative control, two plasmids pAR1219 and pT7-deGFP were introduced in TX-TL system in the absence of IPTG; for a positive control, the same plasmids were used in the presence of IPTG; the aptamer functionality was tested by including all three plasmids at the same concentrations with IPTG induction. The T7 RNAP aptamer was being expressed constitutively since TetR repressor was not present in TX-TL cell extract nor was it encoded on the plasmids used. Eight-fold reduction of deGFP output was obtained in the presence of the plasmid expressing the T7 RNAP aptamer (Supplementary Figure S3C).

After the initial verification of T7 RNAP aptamer activity in TX-TL, the expression cassettes contained in the plasmids pT7-deGFP and pTet-T7apt were joined to create the plasmid pdeGFP-T7apt (Figure 2A). The replication origin and antibiotic resistance was also switched for pdeGFP-T7apt since we aim to investigate its *in vivo* activity. Also, mutant variants of the T7 RNAP aptamer were constructed through site-directed mutagenesis in order to disrupt the conserved sequence domain presumably responsible for T7 RNAP inhibition [37]. The purpose of mutant constructs was two-fold: first, to demonstrate that T7 RNAP inhibition was not mediated by the resource exhaustion caused by T7 RNAP aptamer overproduction [54] and second, to verify the functional importance of conserved sequence domain. The nucleotides 28–31 of the T7apt sequence was targeted and replaced by AAAA, GGGG, or deleted, resulting in three aptamer variants: pdeGFP-T7apt-4A, pdeGFP-T7apt-4G, and pdeGFP-T7apt-4Δ (Figure 2C).

These new set of plasmids were tested together with pAR1219 in the TX-TL expression system (Figure 2B). The results in the TX-TL system were as expected: the correct T7 RNAP aptamer (T7apt) showed 8-fold inhibition of the GFP output as in the previous experiment, whereas the mutant aptamers showed at most 2-fold inhibition, indicating that the core structure of the aptamer was presumably disrupted as a result of mutation. The GFP expression was observed after about 1 hr of TX-TL reaction possibly due to slow production and folding of T7 RNAP, and soon afterwards the slope of GFP production showed clear distinction depending on the identity of input aptamer used. Notably, the GFP output reduction was less than 1.5 fold for T7apt-4Δ; it is plausible that overexpression of non-specific RNA species could cause a small reduction in GFP expression due to resource usage [54].

### Characterization of T7 RNAP aptamer circuit *in vivo*

Encouraged by the activity of T7 RNAP aptamer in the cell-free TX-TL system, we carried out the characterization of circuit *in vivo*, utilizing *E. coli* MGZ1 strain containing constitutively expressed LacI and TetR to complete the inducible circuit. In this circuit, IPTG will induce the production of T7 RNAP, hence driving the expression of deGFP, while aTc will induce the production of T7 RNAP aptamer, repressing the expression of deGFP (Figure 3A). The fluorescence measurement in plate reader clearly showed that aTc-induced T7apt expression inhibited the expression of deGFP (Figure 3B). When T7apt expression was induced with high concentration of aTc (0.6 *µ*g/mL), 5-fold reduction in the deGFP output was obtained compared to no aTc. In contrast, all the mutant variants when induced by the same concentration of aTc showed no effect on the deGFP output (Figure 3B), indicating that the inhibition of T7 RNAP activity is mediated by the correct aptamer rather than by cellular stress due to aTc or overexpression of non-specific RNA species. The fluorescence output decreased once the cells reach stationary phase, however, the fold-change was reasonably consistent over time (Supplementary Figure S4A). Similar results were obtained when *E. coli* were grown in LB or LB with 10 *µ*M IPTG although the foldchange was reduced to 2.5 fold and 2 fold, respectively (Supplementary Figure S4BC).

**Figure 3:**
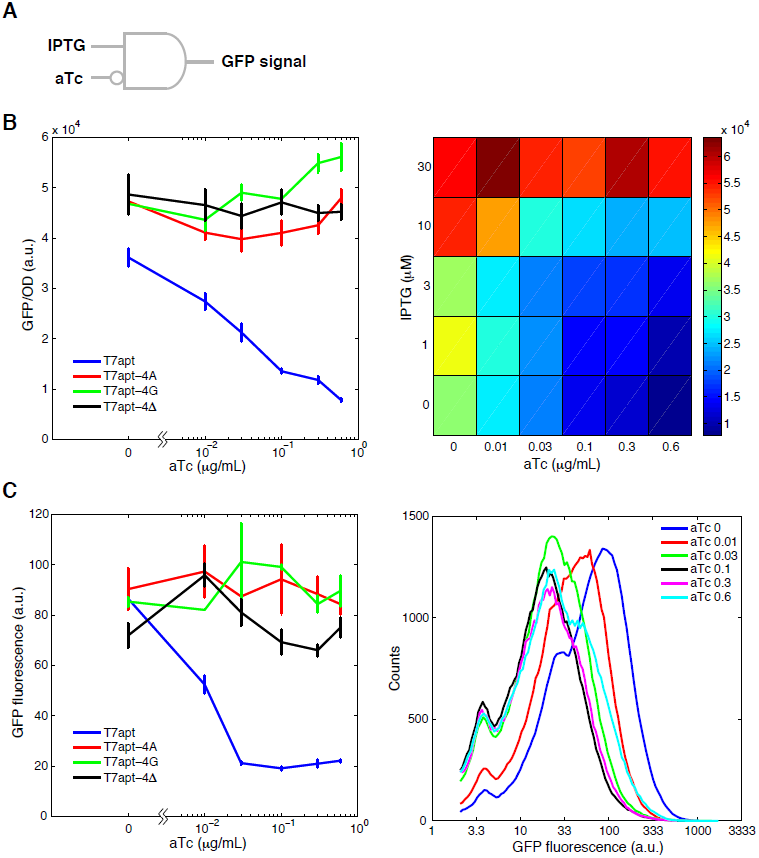
Characterization of T7 RNAP aptamer and its variants *in vivo*. *E. coli* MGZ1 containing constitutively expressed LacI and TetR was used with two plasmids pAR1219 and pdeGFP-T7apt and its variants as shown in Figure 2A to complete the inducible circuit. (**A**) Circuit diagram of T7 RNAP aptamer *in vivo*. Inputs are small molecules IPTG and aTc and the output is GFP fluorescence. (**B**) Plate reader analysis for the response curve of T7 RNAP aptamer and its variants upon aTc induction. GFP/OD values in LB/1% glucose media were measured at 8 hrs for the plot. Time courses are shown in Supplementary Figure S4A. The GFP/OD value heat map for different levels of aTc and IPTG inputs is shown on the right. (**C**) Flow cytometry analysis for the response curve of T7 RNAP aptamer and its variants upon aTc induction. The modal values are plotted with the standard deviation of replicate measurements as error bars. One of the replicate GFP histogram (50,000 events) for T7apt is shown on the right.

A more detailed characterization of the circuit was carried out by testing the response of the circuit to five different IPTG concentrations ranging from 0 to a 30 *µ*M and six different aTc concentrations ranging from 0 to 0.6 *µ*g/mL. As shown in Figure 3B, the response is characterized by high GFP expression for high IPTG concentrations and low aTc concentrations. (The response curves are shown in Supplementary Figure 4D.) At high concentration of IPTG (30 *µ*M), the response was marked by a constant high GFP expression, possibly due to over-expression of T7 RNAP beyond that of T7apt. Even higher concentration of IPTG (100 *µ*M) or aTc (1 *µ*g/mL) severely impaired the cell growth, and therefore, those regions were not further explored. Overall, the circuit exhibited a logic gate behavior as shown in Figure 3A.

To explore the circuit response at the individual cell levels, flow cytometry analysis was performed. Representative flow cytometry histograms of GFP output for different aTc inducer levels clearly showed that GFP output is decreased upon induction of T7apt expression (Figure 3C). The modal fluorescence values from these histograms were used to plot the response curves for T7apt and its variants. The GFP expression showed no dependence on aTc induction levels for mutant T7apt (Figure 3C and Supplementary Figure S5), confirming the findings of plate reader analysis.

## Discussion

Synthetic gene circuitry and cellular programming open up new opportunities for biotechnological and medical applications. With the growing appreciation for RNA molecules as the key component of genetic regulatory networks, efforts to utilize the programmable structure and function of RNA molecules for synthetic biology have accelerated in the recent past. The demonstration of functional RNA aptamer in living cells provides an alternative to simple transcription regulatory circuits with transcription factors, expanding the toolkit to program cells in synthetic biology.

In this work, we show that the T7 RNAP aptamer has the capability of inhibiting the T7 RNAP acivity *in vitro* and *in vivo* with a predictable functional behavior as a modular component within synthetic network. We first demonstrated the activity of T7 RNAP aptamer in *in vitro* transcription circuits. It is not surprising that the aptamer would be functional since the assay environment is similar to the previous works [37]; however, we provide a framework for systematic circuit construction utilizing other RNAP and aptamer components—T3 RNAP and MG aptamer. It remained a question whether T7 RNAP aptamer can show functionality *in vivo* given the complexity of *in vivo* environment with potentially strong crosstalk and degradation for aptamer. We utilized a cell-free TX-TL system as a testbed to quickly assess the feasibility of our approach *in vivo*. Despite certain limitations [54], TX-TL system provides a solid expression platform to connect *in vitro* activity to *in vivo* functionality [38, 52], as recently demonstrated for synthetic RNA circuitry [53]. We used equal concentrations of plasmids for TX-TL experiment to correlate with the plasmids that have pMB1 and p15A origins with similar medium copy numbers—this may partly explain the similar results observed in the aptamer-mediated T7 RNAP repression *in vitro* and *in vivo* (*≈*8-fold vs. *≈*5-fold repression). Together with growing appreciation of the need for testbed in synthetic biology [55, 56, 57], the detailed characterization of cell-free expression platform may provide an easy-to-use breadboard to expedite implementation of synthetic biology circuits *in vivo*. For *in vivo* characterization, *E. coli* MGZ1 cells displayed reasonably high deGFP output even without IPTG induction—this could be due to a weak repression of T7 RNAP expression or a leakage from the medium-copy plasmid, considering that even a small amount of T7 RNAP is sufficient to direct high-level transcription from a T7 promoter. Nevertheless, when cells were induced with IPTG, higher deGFP output was obtained, and repression was observed as expected when induced with aTc (except when IPTG induction was very strong, possibly exceeding the expression level of T7 RNAP aptamer). Utilizing a stronger inducible promoter and improving stability of RNA aptamer using tRNA scaffold [58] or hairpins [59] may increase the dynamic range provided by T7 RNAP aptamer.

A number of straightforward extensions could be implemented for a broad application of the T7 RNAP aptamer. Synthetic circuitry with another aptamer against RNAP, e.g., an aptamer against SP6 RNAP [60], could be utilized to design a synthetic bistable system or oscillators [61, 62]. It is also possible to utilize an antisense signal for T7 RNAP aptamer to produce adaptation behavior [51, 63]. Although widely adapted for gene expression *in vivo*, T7 RNAP due to its very strong transcriptional activity can cause growth arrest and toxicity when strongly induced. Analogous to recent studies utilizing RNA aptamers for quantitation of mRNA [54, 64], concatenation of T7 RNAP aptamer at the 3′ end of an mRNA driven by T7 promoter can provide a necessary negative feedback to regulate the expression levels of target genes, thereby reducing toxicity upon T7 RNAP induction *in vivo*. An intriguing possibility that remains to be tested is whether the T7 RNAP aptamer can provide differential regulation of recently reported synthetic T7 RNAP variants with different promoter specificity [42, 43].

## Supplementary data

Supplementary Data are available

## Funding

This work was supported by National Science Foundation award no. 0832824 (The Molecular Programming Project) and the Defense Advanced Research Projects Agency (DARPA/MTO) Living Foundries program, contract number HR0011-12-C-0065 (DARPA/CMO). The views and conclusions contained in this document are those of the authors and should not be interpreted as representing officially policies, either expressly or implied, of the Defense Advanced Research Projects Agency or the U.S. Government.

## Acknowlegments

The authors thank David Shis and Matthew Bennett for providing the pTara plasmid.

## Conflict of interest statement

None declared.

## Supplementary Information

### Methods

The sequence of DNA oligonucleotides and RNA outputs used for *in vitro* transcription experiments are listed as follows.

#### DNA sequences

pT3-T7apt-nt (77mer), 5′-AAGCAAGGGTAAGATGGAATGAAATTAACCCTCACTAAAGGCGAGCGTAAGTCAATTCCACTATCATTG-CTGCAAGC-3′.

pT3-T7apt-t (49mer), 5′-GCTTGCAGCAATGATAGTGGAATTGACTTACGCTCGCCTTTAGTGAGGG-3′.

pT3-T7apt-v2-nt (83mer), 5′-AAGCAAGGGTAAGATGGAATGAAATTAACCCTCACTAAAGGCGAGCGTAAGTCAATTCCACTATCAT-TGCTGCTTGCCTCGAG-3′.

pT3-T7apt-v2-t (55mer), 5′-CTCGAGGCAAGCAGCAATGATAGTGGAATTGACTTACGCTCGCCTTTAGTGAGGG-3′.

pT3-MGapt-nt (84mer), 5′-AAGCAAGGGTAAGATGGAATGAAATTAACCCTCACTAAAGGACAAGCATCCCGACTGGCGAGAGCCAG-GTAACGAATGGATGCG-3′.

pT3-MGapt-t (56mer), 5′-CGCATCCATTCGTTACCTGGCTCTCGCCAGTCGGGATGCTTGTCCTTTAGTGAGGG-3′.

pT7-MGapt-nt (84mer), 5′-CTAATGAACTACTACTACACACTAATACGACTCACTATAGGACAAGCATCCCGACTGGCGAGAGCCAGG-TAAC-GAATGGATGCG-3′.

pT7-MGapt-t (57mer), 5′-CGCATCCATTCGTTACCTGGCTCTCGCCAGTCGGGATGCTTGTCCTATAGTGAGTCG-3′.

A2-T3 (36mer), 5′-TTAATTTCATTCCATCTTACCCTTGCTTCAATCCGT-3′.

A1-T7 (35mer), 5′-TATTAGTGTGTAGTAGTAGTTCATTAGTGTCGTTC-3′.

#### RNA output sequences

T7apt (38mer), 5′-GGCGAGCGUAAGUCAAUUCCACUAUCAUUGCUGCAAGC-3′.

T7apt-v2 (44mer), 5′-GGCGAGCGUAAGUCAAUUCCACUAUCAUUGCUGCUUGCCUCGAG-3′.

MGapt (45mer), 5′-GGACAAGCAUCCCGACUGGCGAGAGCCAGGUAACGAAUGGAUGCG-3′.

The sequence of DNA oligonucleotides used for cloning and Gibson assembly are listed as follows.

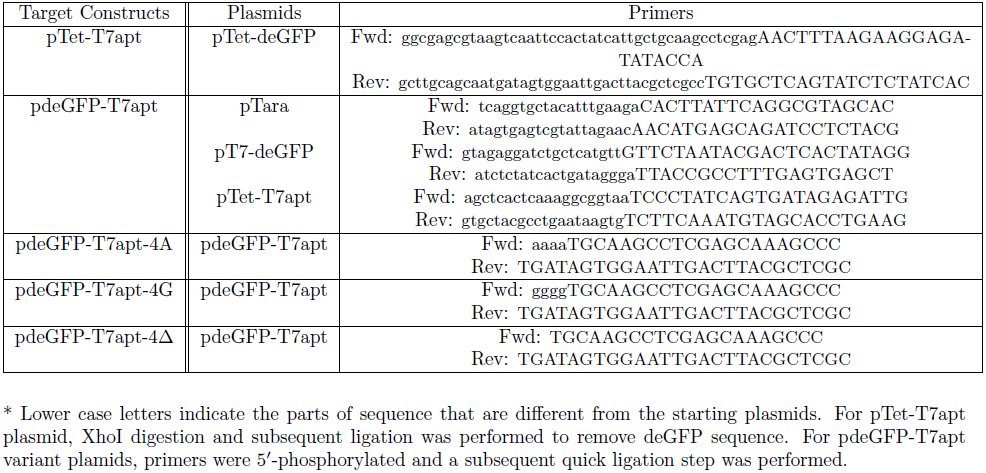

**Figure S1.**
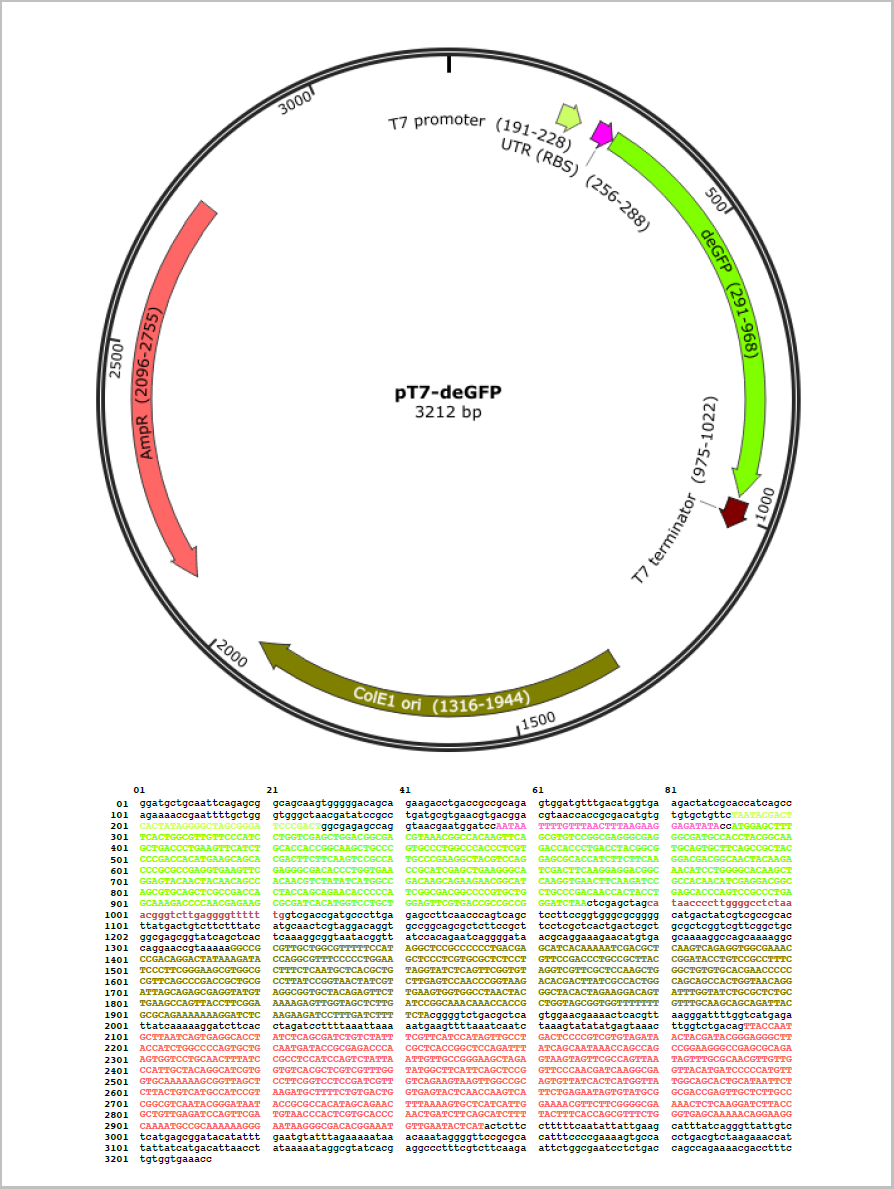
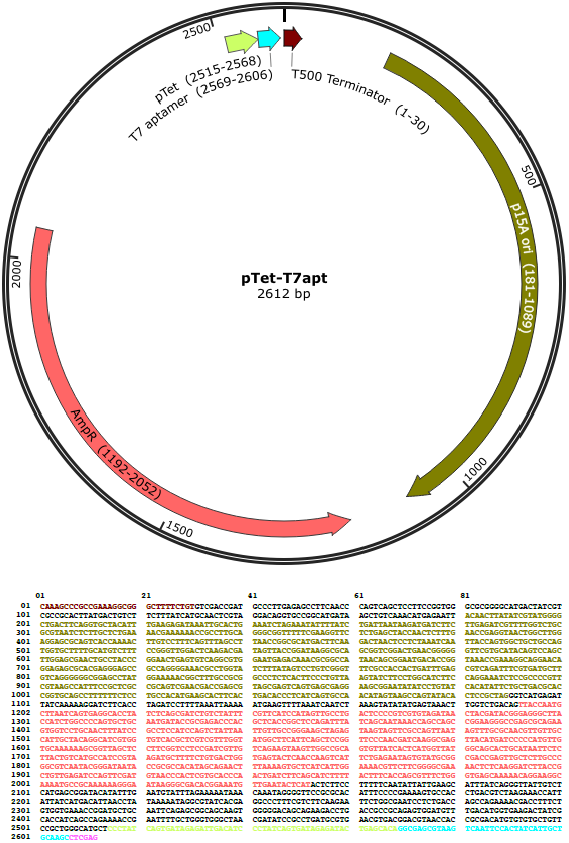
Plasmid maps. Plasmid pT7-deGFP was derived from the plasmid pIVEX2.3d (Roche) [38]. Plasmid pTet-T7apt was constructed using Gibson assembly method with plasmid pBEST-Luc (Promega) as a backbone. Plasmid pTara was used as a backbone to join pT7-deGFP and pTet-aptamer expression cassettes in plasmid pdeGFP-T7apt.

**Figure S2.**
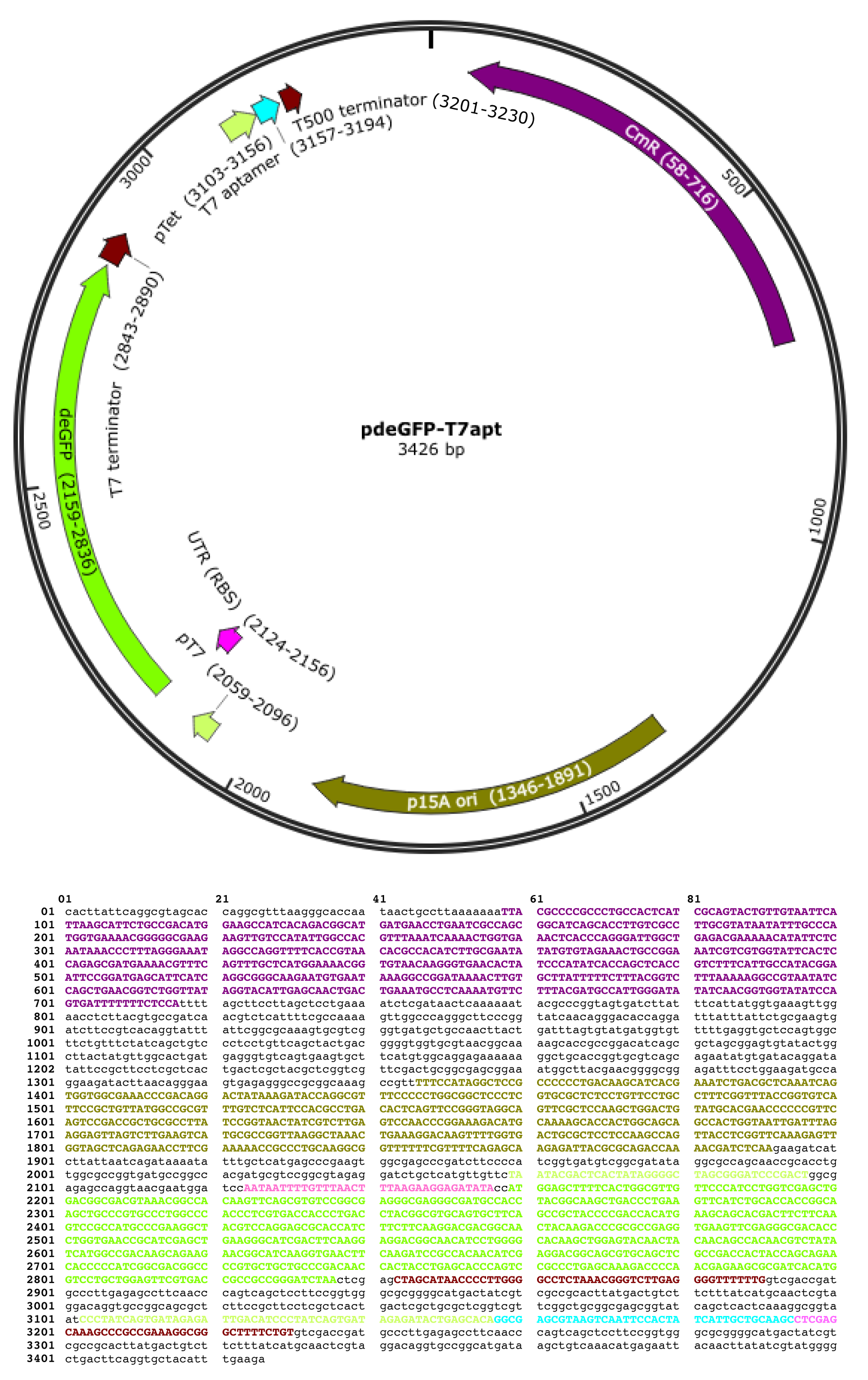
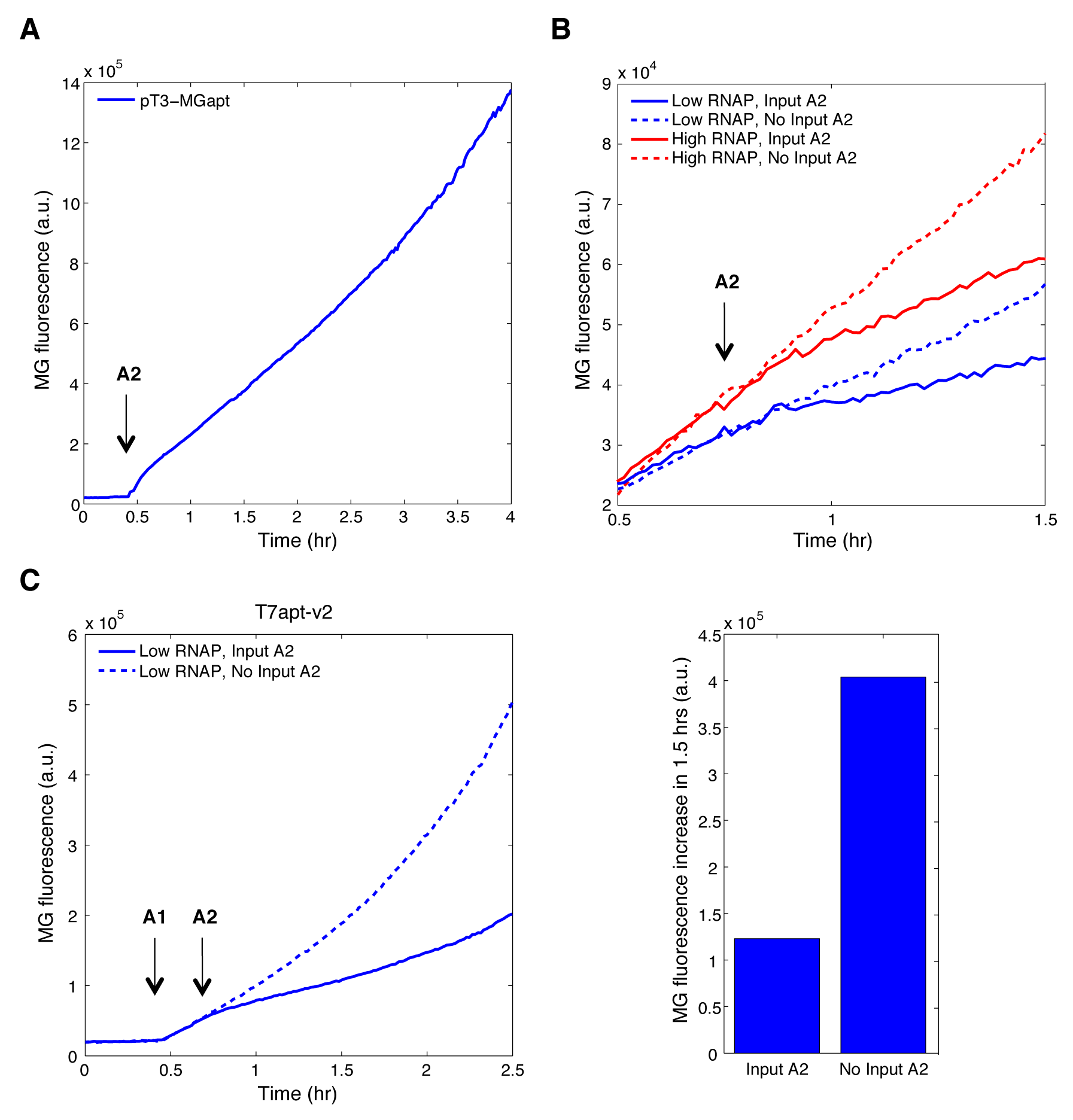
Spectrofluorometer analysis of T7 RNAP aptamer in *in vitro* transcription circuits. (**A**) With pT3-MGapt template in a transcription buffer in a cuvette, T3 RNAP was added at 10 min. MG aptamer was not produced until the activator A2 was added at 25 min, indicating that the incomplete promoter design also works for T3 promoter much as T7 promoter design used for genelet circuits. (**B**) Close-up of the timecourse experiment shown in Figure 1C. The activator A2 was added at 40 min. The ‘input A2’ trajectories began to diverge from ‘no input A2’ trajectories about 10 min after the introduction of input A2. (**C**) Test of modified aptamer against T7 RNAP (T7apt-v2). Here, pT3-T7apt-v2 template was used instead of pT3-T7apt. The two RNA polymerases were added at 10 min, the input A1 was added at 25 min, and the input A2 was added at 45 min following the protocols used in Figure 1C. The T7apt-v2 was efficient in repressing the T7 RNAP activity; about 3-fold decrease in MGapt signal was observed over 1.5 hours in the presence of the T7apt-v2.

**Figure S3.**
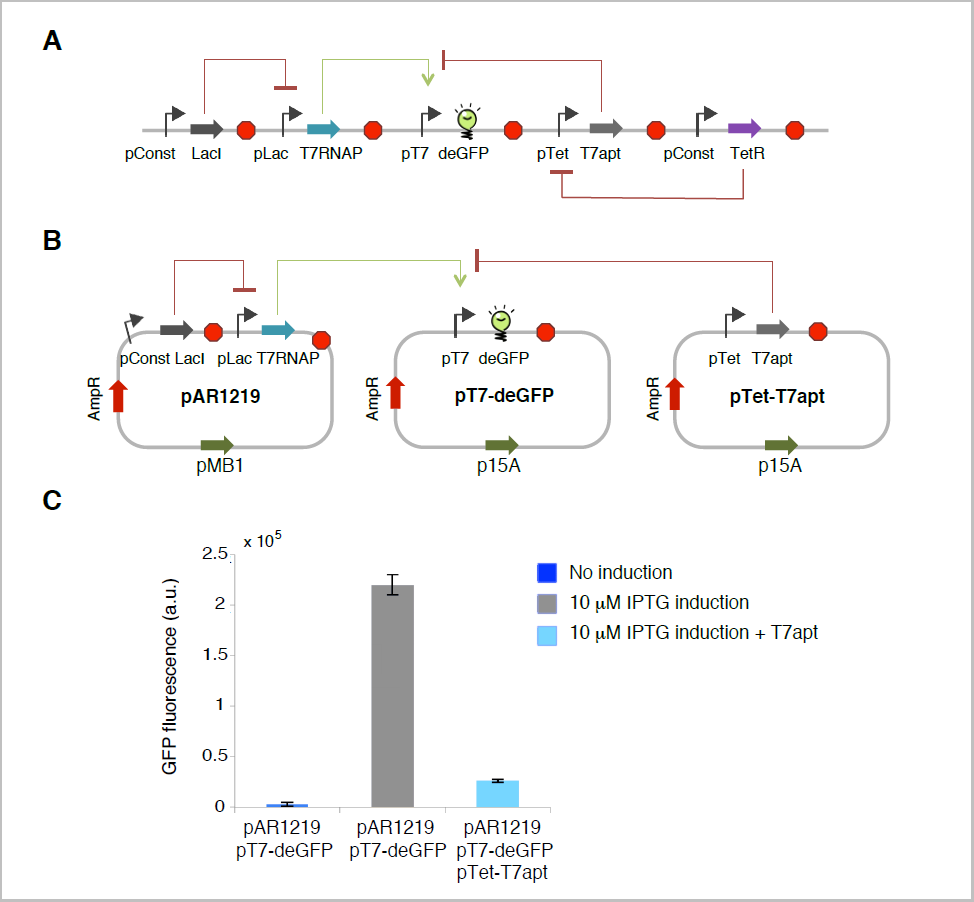
Genetic circuits and aptamer-mediated T7 RNAP inhibition in the TX-TL system. (**A**) Designed circuit for *in vivo* experiments. (**B**) Three-plasmid system used for aptamer-mediated T7 RNAP inhibition test in the TX-TL cell-free extract. (**C**) Experimental results showed that T7 RNAP was induced by IPTG to express deGFP and that deGFP expression was decreased by 8 fold in the presence of T7 RNAP aptamer expressing plasmid pTet-T7apt. The T7 RNAP aptamer production was not inhibited by TetR since the cell extract does not contain TetR protein. The fluorescence signal from deGFP were measured at 4 hrs.

**Figure S4.**
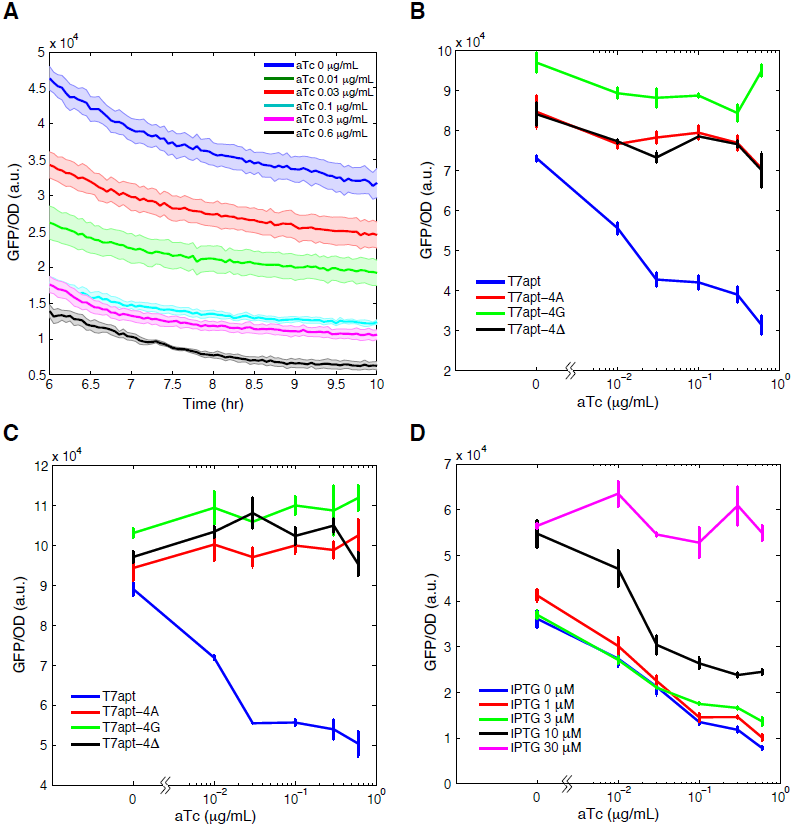
Plate reader analysis of T7 RNAP aptamer and variants *in vivo*. (**A**) Time course of GFP/OD measurements in LB/1% glucose media as reported in Figure 3B left panel. The GFP/OD values decreased over time as the cells reached stationary phase after 6 hrs. However, the relative ratio of GFP/OD values for different aTc induction levels did not vary considerably during the 6 to 10 hr time window. (**B**) The response curve of T7 RNAP aptamer and its variants upon aTc induction in LB media. (**C**) The response curve of T7 RNAP aptamer and its variants upon aTc induction in LB media with 10 *µ*M IPTG. (**D**) The response curve of T7 RNAP aptamer upon aTc induction in LB/1% glucose media with different levels of IPTG. At high IPTG level, the response to aTc induction was abolished. This set of data is represented as a heat map in Figure 3B right panel. GFP/OD values were measured after 8 hrs for (B–D).

**Figure S5.**
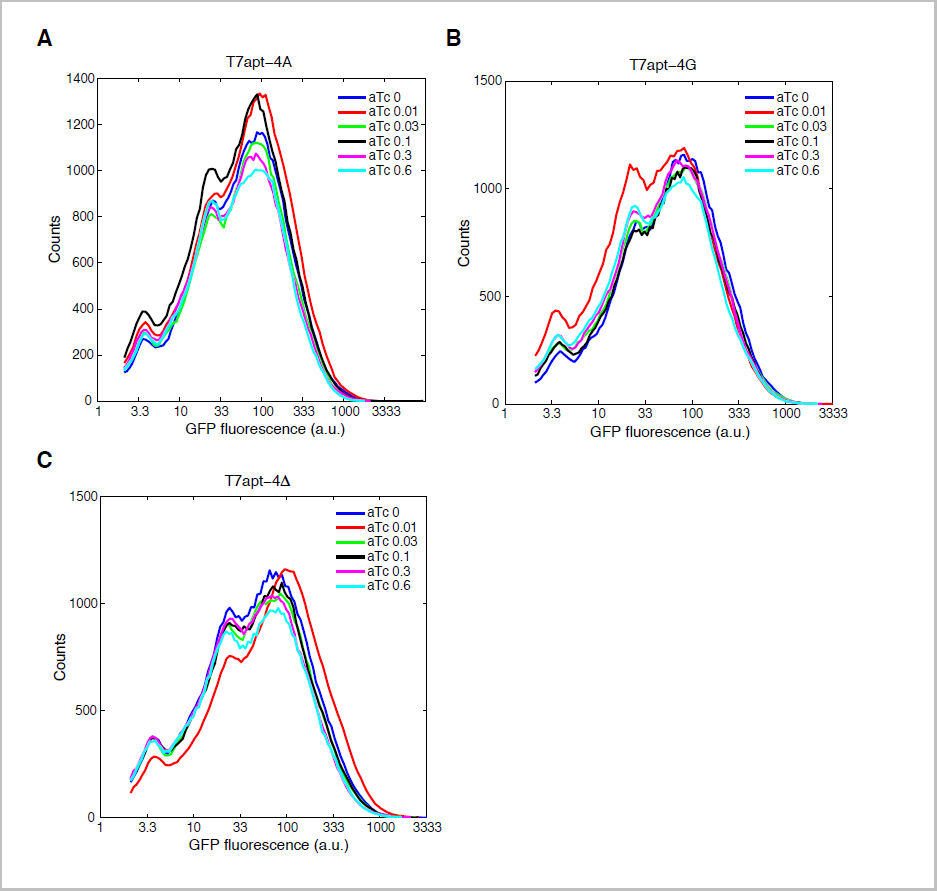
Flow cytometry analysis of mutant T7 RNAP aptamer *in vivo*. One of the replicate GFP histogram (50,000 events) for (**A**) T7apt-4A, (**B**) T7apt-4G, (**C**) T7apt-4Δ. The GFP expression levels did not change upon induction of T7 RNAP aptamer by aTc for all mutant aptamers. The modal GFP values are plotted in Figure 3C.

